# A tissue-specific self-interacting chromatin domain forms independently of enhancer-promoter interactions

**DOI:** 10.1101/234427

**Authors:** Jill M Brown, Nigel A Roberts, Bryony Graham, Dominic Waithe, Christoffer Lagerholm, Jelena M Telenius, Sara De Ornellas, A Marieke Oudelaar, Izabela Szczerbal, Christian Babbs, Mira T Kassouf, Jim R Hughes, Douglas R Higgs, Veronica J Buckle

## Abstract

A variety of self-interacting domains, defined at different levels of resolution, have been described in mammalian genomes. These include Chromatin Compartments (A and B)^1^, Topologically Associated Domains (TADs)^2,3^, contact domains^4,5^, sub-TADs^6^, insulated neighbourhoods^7^ and frequently interacting regions (FIREs)^8^. Whereas many studies have found the organisation of self-interacting domains to be conserved across cell types^3^^8^^9^, some do form in a lineage-specific manner^6,7,10^. However, it is not clear to what degree such tissue-specific structures result from processes related to gene activity such as enhancer-promoter interactions or whether they form earlier during lineage commitment and are therefore likely to be prerequisite for promoting gene expression. To examine these models of genome organisation in detail, we used a combination of high-resolution chromosome conformation capture, a newly-developed form of quantitative fluorescence *in-situ* hybridisation and super-resolution imaging to study a 70 kb self-interacting domain containing the mouse α-globin locus. To understand how this self-interacting domain is established, we studied the region when the genes are inactive and during erythroid differentiation when the genes are progressively switched on. In contrast to many current models of long-range gene regulation, we show that an erythroid-specific, decompacted self-interacting domain, delimited by convergent CTCF/cohesin binding sites, forms prior to the onset of robust gene expression. Using previously established mouse models we show that formation of the self-interacting domain does not rely on interactions between the α-globin genes and their enhancers. As there are also no tissue-specific changes in CTCF binding, then formation of the domain may simply depend on the presence of activated lineage-specific cis-elements driving a transcription-independent mechanism for opening chromatin throughout the 70 kb region to create a permissive environment for gene expression. These findings are consistent with a model of loop-extrusion in which all segments of chromatin, within a region delimited by CTCF boundary elements, can contact each other. Our findings suggest that activation of tissue-specific element(s)within such a self-interacting region is sufficient to influence all chromatin within the domain.

---

The mouse α-globin cluster is contained within a well-characterised 70kb self-interacting domain in which we have previously identified all *cis*-acting elements, including promoters, enhancers and CTCF/cohesin binding sites (Fig. 1)^11^. In mES cells the α-globin promoters and five enhancers are not bound by transcription factors and the genes are silent^12^. Further, we detect no evidence of a strong self-interacting domain in mES cells, whereas such a structure is clearly present in differentiating erythroblasts^10^. Yet the largely convergent boundary elements are occupied by CTCF and cohesin in both cell types^13^, suggesting that CTCF/cohesin are not the primary mediators of this tissue-specific domain formation. To determine the relationship between activation of the α-globin gene cluster and formation of a self-interacting domain, we examined fetal liver cells from E12.5 mice (MFL), cultured and harvested *ex-vivo* at two stages of differentiation. Initially (MFL 0h) the cells correspond to proerythroblasts in which all *cis*-elements are bound by transcription factors but there is little or no α-globin expression. After 30 hours (MFL 30h), corresponding to intermediate erythroblasts, the α-globin genes are fully active and transcribed at maximal levels (Fig. 1a, b). NG Capture-C analysis of chromatin interaction frequency^10^ at the α-globin locus in MFL 0h and 30h populations indicates that the α-globin self-interacting domain is present and apparently equivalent in both populations (Fig. 1c). NG Capture-C analysis from one of the flanking CTCF/cohesin binding sites (-39.5) at the border of the domain shows that this site does not interact with the enhancers and promoters within the self-interacting domain but rather with the convergent CTCF/cohesin binding sites on the opposite side of the domain, at both 0h and 30h time points. Concordant with evidence in neural development^14^, we conclude that the α-globin self-interacting domain, which is absent in mouse ES cells^10^ (Extended Data Fig. 1), is already formed at an early stage of erythroid differentiation, prior to the onset of robust α-globin transcription.

**Figure 1:**
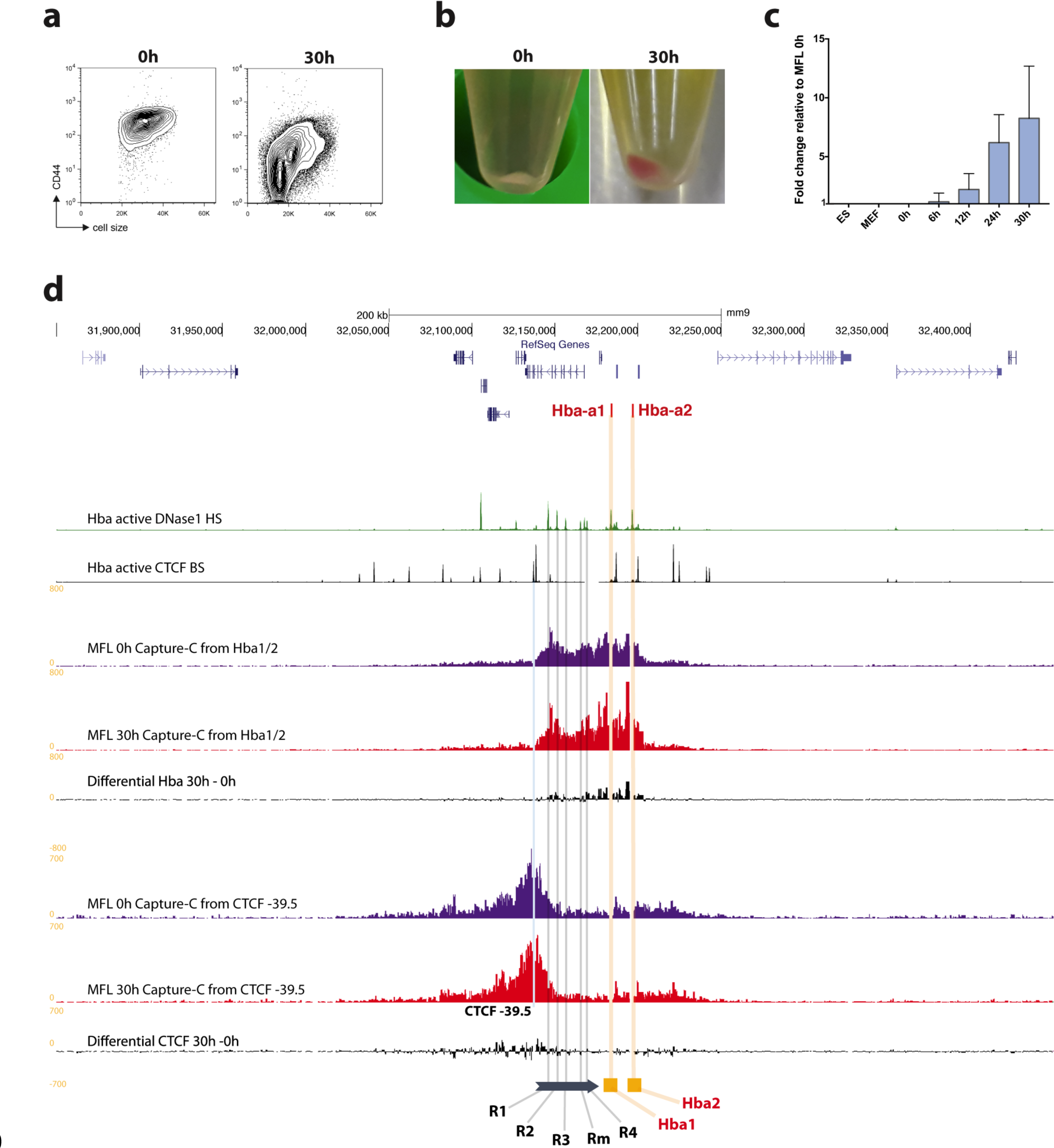
A self-interacting domain at the α-globin locus is formed in both MFL 0h and 30h erythroblasts. **a,** Representative FACS plots of MFL erythroblasts defined by CD44 and cell size, at 0h and after a further 30h differentiation *in vitro,* identifying distinct populations at the two timepoints. **b,** Cell pellets from MFL 0h and 30h culture showing differentiated haemoglobinised erythroblasts present only at 30h. **c,** Nascent Hba transcription relative to 18S in the cell types and MFL timepoints indicated. n=3. Error bar is standard deviation. **d,** Map of the gene dense murine α-globin locus with Hba genes highlighted in red and gene browser tracks showing DNase1 hypersensitive sites (green) and CTCF binding sites (BS black)^11^. Next, NG Capture-C tracks using Hba1/2 as viewpoints in MFL erythroblasts at 0h and 30h with a differential track showing minimal changes between the two timepoints. Three further tracks in the same arrangement use the CTCF BS −39.5 as viewpoint. The location of the Hba genes, the five murine enhancer elements and the CTCF BS −39.5 are marked against the browser tracks in yellow, grey and blue vertical bars respectively.

Rather than discrete contacts between elements, NG Capture-C has demonstrated that the α-globin self-interacting domain reflects interactions between all sequences within a 70kb region of chromatin lying between CTCF boundaries^10,11^. This generally increased contact frequency could reflect a spatially confined volume or alternatively a more dynamically interactive region of chromatin. As 3C data only quantify the ligations between juxtaposed chromatin segments in a population of cells, they do not distinguish between these two possibilities. Guided by the 3C interaction frequency data, we positioned panels of FISH probes across the α-globin locus, allowing us to determine the changes in 3D structure of this locus during erythroid differentiation at single cell level, and the roles that key cis-sequences play in forming the self-interacting domain.

For FISH probes, we used a BAC (COMP) precisely covering the α-globin self-interacting domain, and two BACs (F1 and F2) flanking this region (Fig. 2). We also designed smaller (~7kb) plasmid probes (Ex, E, A, C and Cx) to analyse chromatin organisation in finer detail. The Anchor ‘A’ probe was located at the distal extremity of the domain and is the nearest region of unique sequence adjacent to the α-globin genes. All measurements involving the plasmid probes were made relative to this position. ‘Ex’ defines the proximal edge of the domain and ‘E’ was sited at the two major enhancer elements MCS-R1 and MCS–R2. Two control probes (‘C’ and ‘Cx’) were positioned outside of the self-interacting domain, in a region showing little interaction by NG Capture-C with the α-globin genes or their enhancers. These control probes are equidistant in linear sequence to A as upstream probes E and Ex respectively. Prior to the FISH experiments, we performed NG Capture-C on MFL 30h erythroblasts and mES cells using capture oligonucleotides corresponding to the central points of each plasmid probe, to determine the interactions they detect across the locus (Extended Data Fig. 1). FISH with this probe panel then allowed us to measure 3D distances and volumes occupied by the chromatin within and outside the α-globin domain in single cells.

**Figure 2:**
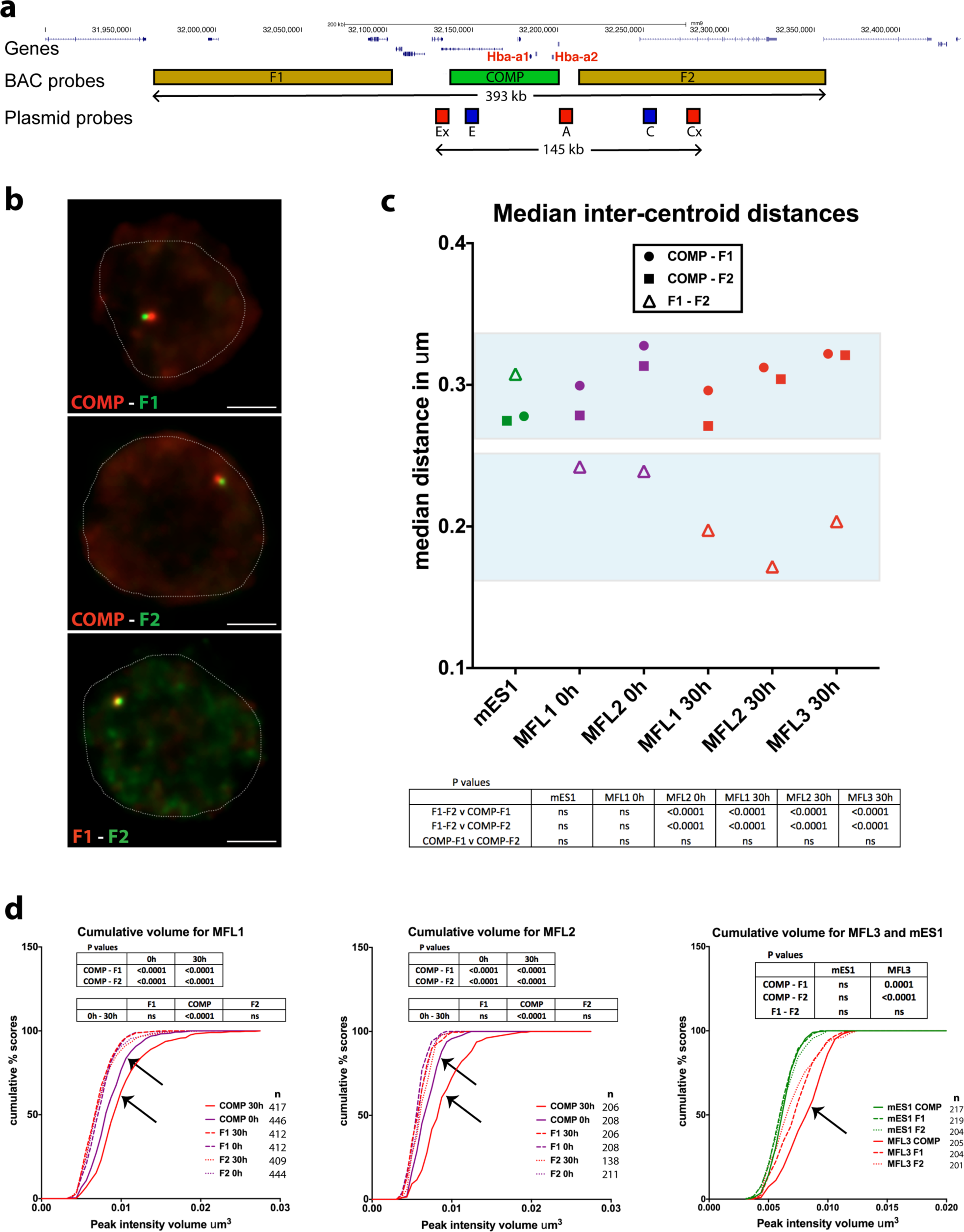
Volume and proximity measurements across the α-globin locus in WT mice. **a,** Gene map and locations of the BAC and plasmid FISH probes to scale, with total genomic distance encompassed. **b,** Representative images for the three BAC probe pairs in MFL 30h erythroblasts with nuclei delineated. Scale bar 2 μ™. **c,** Median inter-centroid distances between the three probe pairs indicated, in mES cells (green) and erythroblasts at 0h (purple) and 30h (red) timepoints. MFL1, 2 and 3 represent cultures from three individual foetal livers. Light blue boxes emphasise proximity of the flanking regions F1-F2 in erythroblasts. n = 87-236 - see Extended Data Fig. 2 for the complete data set. **d,** Cumulative frequency plots of BAC signal volumes in mES cells (green) and erythroblasts at 0h (purple) and 30h (red). COMP values indicating expanded volume are arrowed. All P values are derived by a Kruskal-Wallis test with Dunn’s multiple comparisons. ns = not significant.

To ensure that we preserved nuclear structure of the cells being analysed, we developed a method Raser-FISH (Resolution After Single-strand Exonuclease Resection) for labeling chromosomal loci without denaturing the DNA (see Methods). Using COMP, F1 and F2, we used this approach to measure inter-probe distances in mES cells and MFL 0h and 30h erythroblasts with three paired combinations (COMP-F1, COMP-F2 and F1-F2) (Fig. 2a, b). We found no significant differences between COMP-F1 or COMP-F2 in mES cells, MFL 0h and MFL 30h erythroblasts (Fig. 2c and Extended Data Fig. 2). However, we noted that although the linear genomic distance between F1 and F2 is twice that of COMP-F1 or COMP-F2, the median inter-probe distances between F1 and F2 are nevertheless shorter at MFL 0h (Fig. 2c). Importantly at MFL 30h, we observed a further, highly significant shortening of F1-F2 measurements compared to COMP-F1 and COMP-F2 (Fig. 2c). By contrast, there is no significant difference between distance measurements for the three probe pairs in mES cells. To analyse this further, we calculated the Pearson’s correlation for the BAC signal pairs to estimate the degree of signal overlap. There is marked overlap between F1 and F2 signals at 0h (median coefficients 0.6-0.61) and further overlap at 30h (median coefficients 0.66-0.75) in contrast to mES cells (median coefficient 0.43). Thus it appears that as erythroblasts differentiate and transcription is up-regulated, the flanking regions of the self-interacting domain are more frequently found in close proximity.

Volume measurements of probe signals can distinguish between a compact structure and a decompacted, dynamically interacting region. The genomic length of the COMP probe is 64 kb compared to 139 kb for both F1 and F2. The volume of the COMP BAC signal was greater at MFL 0h than F1 or F2 and the difference increased at MFL 30h. By contrast in mES cells, the three sets of volume measurements were comparable (Fig. 2d). This indicates that chromatin within the domain becomes less compact during differentiation relative to surrounding chromatin by MFL 0h and decompacts further during subsequent erythroid differentiation to consolidate the three-dimensional distinction of this self-interacting domain.

To further test these observations, we used the precisely positioned pairs of plasmid probes (A-E, A-C, A-Ex, and A-Cx), to analyse intra-chromosomal distances in mES and erythroid cells (Extended Data Fig. 3, 4). As for the BAC probes, we found no significant differences in mES cells between measurements across the self-interacting domain versus the control region (A-Ex versus A-Cx). This finding is supported by NG Capture-C data from the viewpoints of all five plasmid probes where only immediate proximity interactions are detected (Extended Data Fig. 1), indicating that no domain of interaction is present at this site in mES cells. Analysing early erythroblasts (MFL 0h), although we find no difference in the median measurements between A-E and A-C, we do find significant difference between A-Ex and A-Cx measurements, in agreement with NG Capture-C data (Fig. 1) where interaction frequencies indicate that the α-globin domain has already formed. In intermediate erythroblasts (MFL 30h) when the α-globin genes are fully active, the difference becomes highly significant for A-Ex versus A-Cx, and with A more frequently closer to Ex and to E, whilst distances between A to C and to Cx increase. These data indicate that even when the α-globin genes are silent or minimally active, proximity has already been established between the extremities of the self-interacting domain. In intermediate erythroblasts when >80% of the α-genes are highly active, the extremities are yet more frequently in close proximity, supporting the concept of a dynamically looped domain where sites A and Ex sit in regions defining the borders of generalised interactions. Importantly, the persistence of the spread of the distance measurements observed from individual loci throughout differentiation (Extended Data Figs. 1 and 4) implies that this self-interacting domain is not a static loop but rather a dynamic structure that exists in different conformations at any one time point, in agreement with a proposed model based on CTCF/cohesin complex dynamics^15^.

To visualize this region at the highest possible resolution we analysed erythroblasts using Stimulated Emission Depletion (STED) imaging^16^. As probes, we used the COMP BAC, which precisely corresponds to the self-interacting domain and the two flanking plasmid probes (A and Ex), detecting the extremities of the domain (Fig. 3a). In 24 out of 35 erythroblast nuclei, the signals for A and Ex were juxtaposed or overlapping, with the COMP signal distinctly to the side or wrapped around them (Fig. 3b).

**Figure 3:**
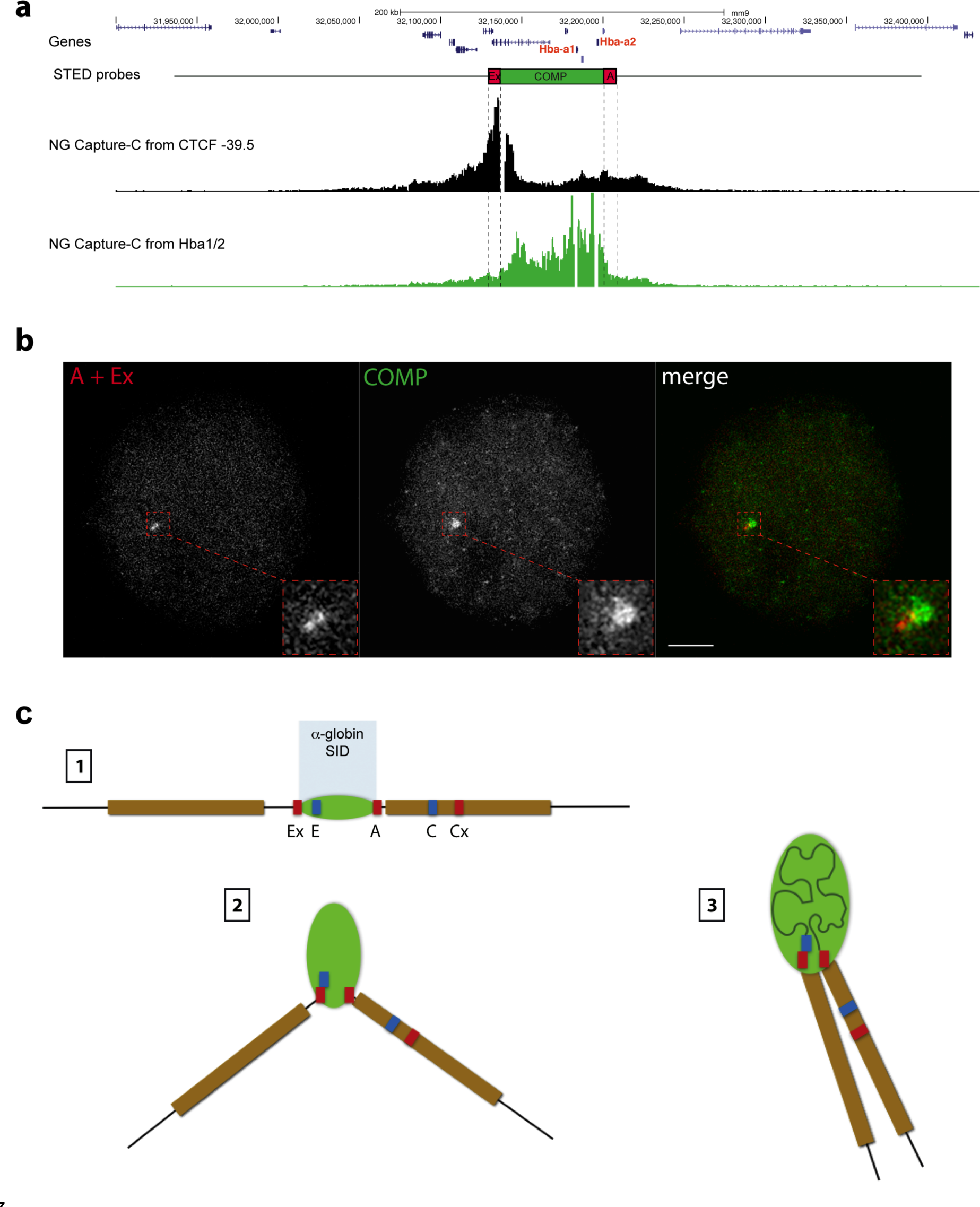
Super-resolution imaging of the α-globin domain informs a schematic model of the locus. **a,** Gene map with FISH probe locations marked against NG Capture-C tracks depicting interactions at MFL 30h from the viewpoints CTCF BS −39.5 (black) and HBA1/2 promoters (green). **b,** 2D STED maximum intensity images of FISH probes A and Ex (both red) which flank the α-globin domain, the COMP BAC (green) defining the extent of the domain and a merged image showing a cloud of domain signal distinct from the paired probes A and Ex. Bar = 2 μm. **c,** Model showing the development of the α-globin self-interacting domain (SID) (green). Sites detected by FISH probes are as for Fig. 2. Stage 1 represents the linear locus, whilst Stage 2 and 3 depict the development of the self-interacting domain, where the domain expands as chromatin decompacts and the flanking regions can sit in proximity.

Based on all the data above, we present a model (Fig. 3c) for the formation of an active regulatory domain at an early stage of erythropoiesis, with the self-interacting domain boundaries and external flanking chromatin frequently sited closer together and chromatin within the domain becoming less compact relative to surrounding regions. These features become more pronounced over the course of erythroid differentiation. Notably, NG Capture-C from the control sites C and Cx shows their avoidance of the α-globin domain but does detect infrequent interactions between the flanking chromatin regions (Extended Data Fig. 5), suggesting low frequency contacts of surrounding chromatin caused by formation of the domain.

It has been proposed that self-interacting domains might result from specific interactions between enhancers, promoters and CTCF/cohesin elements^17-19^. Recent data^13^, together with the evidence presented here, show that rather than interacting directly with the α-globin enhancers and promoters, the flanking CTCF sites appear to avoid these elements: in fact, each shows interaction with the opposite flanking regions. NG Capture-C analysis indicates that the self-interacting domain still forms in engineered mice with a homozygous double knockout of the two major α-globin enhancers (DKO for MCS-R1 and MCS-R2), although interactions within the domain appear somewhat reduced^20^. Here we examined inter-probe distances in erythroblasts from such mice in which nascent transcription from the α-globin locus is reduced by 90%. This showed that a self-interacting domain still forms in the absence of the major enhancers (Fig. 4a). Notably we found a highly significant overlap of F1 with F2 signals calculated by Pearson’s correlation (coefficients WT 0.75 and DKO 0.67) when compared to COMP with F1 or F2 (coefficients WT 0.46 and DKO 0.46). For plasmid hybridisations, there was also a significant difference in distance measurements for both A-E versus A-C and A-Ex versus A-Cx. (Fig. 4c, Extended Data Fig. 6a, b). This indicates that the structure we detect in WT erythroblasts, where the extremities of the domain come together, is recapitulated in the double enhancer knockout despite the physical absence of the core binding sites of the two enhancers and the severe reduction in transcriptional output from the α-globin genes. We next analysed erythroblasts from a mouse homozygous for a 16kb deletion that removes both α-globin genes from each chromosome (AMKO), consequently there is no adult α-globin transcription at all. As for WT, we found a significant difference in measurements within and outside of the self-interacting domain (Fig. 4d, Extended Data Fig. 6c) and this is matched by NG Capture-C analysis, which still detects a definable self-interacting domain in the absence of the α-globin genes (Fig. 4e). Here again we see evidence that the domain structure still forms, this time in the absence of the α-globin promoters and of transcription from the α-globin genes.

**Figure 4:**
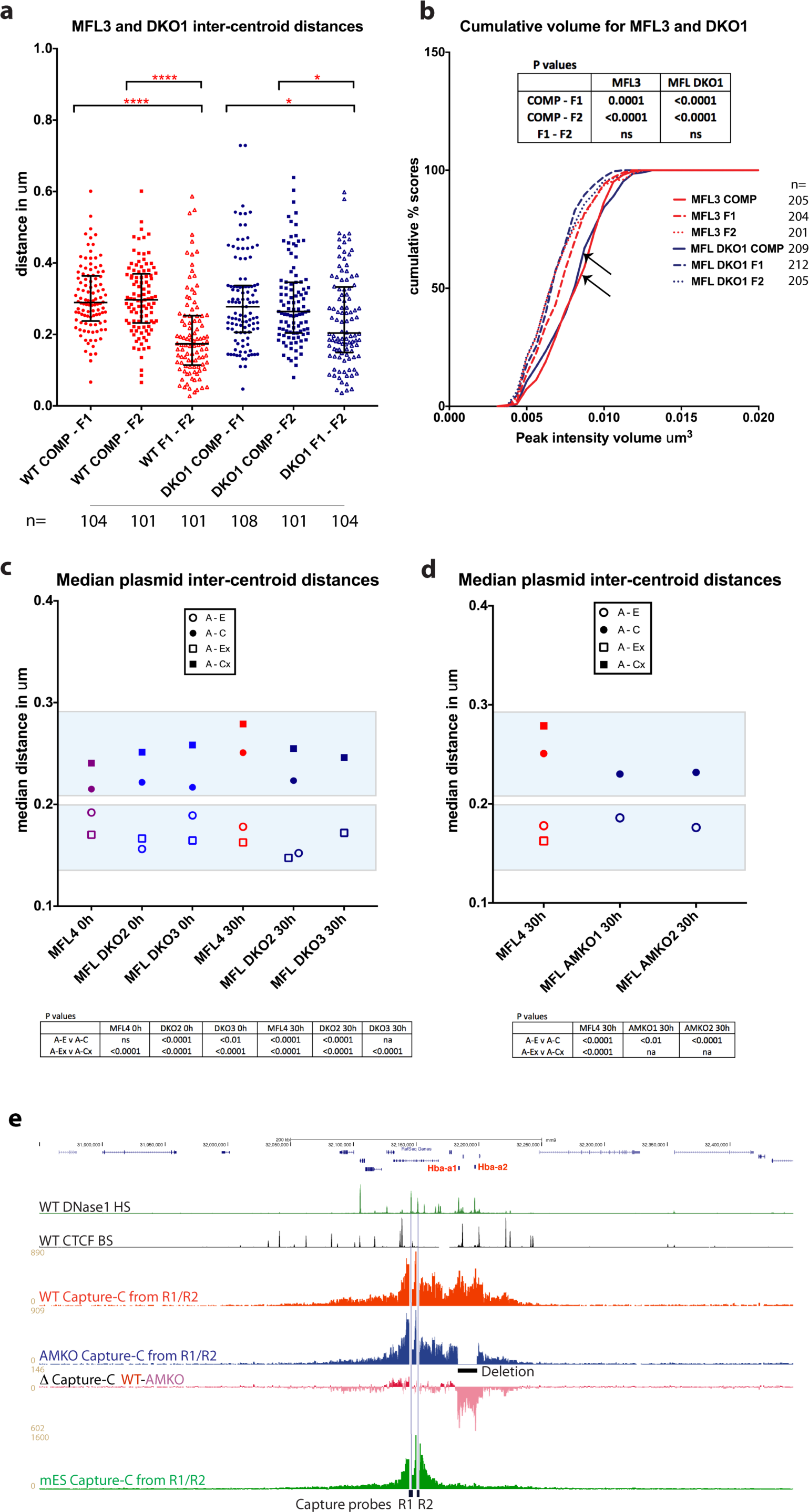
The α -globin domain still forms in the absence of critical elements. **a,** Pairwise intercentroid distances between three BAC probes in 30h erythroblasts derived from littermates MFL3 (WT) and DKO1 (homozygous deletions for MCSR1/R2). F1-F2 are significantly more frequently closer than COMP-F1 and COMP-F2 in both WT (p<0.0001 for both) and DKO1 derived erythroblasts (p=0.0374 and 0.0468 respectively). **b,** Cumulative frequency plots of BAC signal volumes in 30h erythroblasts from MFL3 and DKO1. COMP values are arrowed. **c,** Median inter-centroid distances between four plasmid probe pairs at MFL 0h and 30h from littermates WT MFL4 and two homozygous double knockout embryos DKO2 and DKO3. Light blue boxes emphasise the shorter distances within the self-interacting domain in both WT and knockouts. See Extended Data Fig. 6 for the complete data set. **d,** Median inter-centroid distances between plasmid probe pairs A-Ex (represented as A-E distance because of a 16 kb α-globin gene deletion) and A-C at MFL 30h in two α-globin knockout lines from littermates AMKO1 and AMKO2, plotted against WT MFL4. Light blue boxes are as for c. See Extended Data Fig. 6 for full data. **e,** Gene map followed by DNase1 hypersensitive sites (green) and CTCF BS (black) genome browser tracks, then NG Capture-C tracks highlighting interactions from MCS-R1/-R2 viewpoints in MFL WT (red) and MFL AMKO (blue), with a differential track WT-AMKO showing persistence of domain structure in AMKO when contrasted with the absence of a domain observed in mES cells (green). All P values are derived by a Kruskal-Wallis test with Dunn’s multiple comparisons. ns = not significant.

Finally we asked whether transcriptional up-regulation at the α-globin locus is directly related to chromatin decompaction within the self-interacting domain. Volume measurements of the FISH signal generated using the COMP probe, together with the two flanking probes F1 and F2, indicate that the chromatin within the domain is decompacted compared to the flanking regions in both wild-type and DKO knockout cells (Fig. 4b). Hence, within the domain, chromatin decompaction is uncoupled from transcriptional upregulation of the α-globin genes, indicating a response to earlier events at the locus. Decompaction of chromatin has also been uncoupled from enhancer-promoter interaction at the Shh locus in mice^21^.

In contrast to many current models of long-range gene regulation, we have shown, using a combination of NG Capture-C, quantitative FISH and super-resolution microscopy, that an erythroid specific decompacted self-interacting domain, delimited by convergent CTCF/cohesin binding sites forms prior to gene up-regulation and does not rely on interactions between the α-globin genes and their enhancers or detectable tissue-specific changes in CTCF binding^13^. Our findings therefore suggest that formation of the domain more simply depends on the presence of activated lineage-specific *cis*-elements driving a transcription-independent mechanism for opening chromatin. Certainly dCas9 gene activation alone has been shown to be insufficient to create a domain structure^14^, and in Drosophila more generally, TAD formation can arise independently of transcription^22^. Our findings could be explained by the recently proposed mechanism of chromatin loop extrusion^23-25^. This is thought to be an active process in which a loop-extruding factor, containing two DNA-encompassing units such as cohesin, associates with chromatin and travels along the chromatin fibre in opposite directions, creating a progressively larger intervening loop, until the factor is stalled at appropriately orientated CTCF-bound elements. This model would explain our observations that CTCF/cohesin sites flanking the α-globin self-interacting domain become juxtaposed around a decompacted loop of chromatin in erythroblasts. In this model, domains could arise from a dynamic balance of cohesin loading and removal and loop extrusion and blocking^26^. We have previously noted an accumulation of cohesin in erythroid cells around all five enhancer-like elements of the α-globin cluster^13^, which may act as entry points for cohesin. In this study, the remaining erythroid-specific elements in the absence of MCS-R1, MCS-R2 and α-globin genes are MCS-R3, MCS-R4 and Rm, which could play a redundant role in the formation of the self-interacting domain. This would explain a role for *cis*-elements like MCS-R3 and –Rm, which have the signature of enhancers but without obvious enhancer activity^20^, that is distinct from, and is active prior to, gene up-regulation. In this scenario and compatible with our data, the boundary elements of the domain are dynamically brought into proximity as a result of loop extrusion or similar mechanism, rather than initiating the formation of a self-interacting domain.

## Methods

### Animal Procedure

The mutant and wild type mouse strains reported in this study were maintained on a mixed background and were generated and phenotyped in accordance with Animal [Scientific Procedures] Act 1986, with procedures reviewed by the clinical medicine Animal Welfare and Ethical Review Body (AWERB), and conducted under project licence PPL 30/3339. All animals were singly housed, provided with food and water *ad libitum* and maintained on a 12 h light: 12 h dark cycle (150–200 lux cool white LED light, measured at the cage floor). Mice were given neutral identifiers and analysed by research technicians unaware of mouse genotype during outcome assessment.

### Cell Culture

Erythropoiesis can be faithfully recapitulated *ex vivo* where progenitor cells differentiate down an erythroid pathway, making all necessary proteins for red cell function, to a late stage when the nucleus condenses and is finally extruded from the cell. *Ex vivo* culture of foetal liver cells from E12.5 mice (MFL) allows us to access erythroblasts at an early stage of differentiation with low levels of globin transcription (MFL 0h) and at an intermediate stage when erythroblasts are at peak transcription of the globin genes (MFL 30h). Previous analysis of chromatin conformation at the *α-globin* locus has used Ter119-positive erythroblasts derived from adult spleen to represent the *α-globin-on* population^10,13^. Data derived from these cells and from MFL 30h erythroblast cultures are comparable (Extended Data Fig. 7). Our *in vitro* mouse foetal liver (MFL) culturing system is based on previous protocols^27,28^. Briefly, MFL cells, taken at E12.5, were cultured in StemPro medium (Invitrogen) supplemented with Epo (1 U/mL) (Janssen, PL 00242/029), SCF (50 ng/mL) (Peprotech, 250-03), dexamethasone (μM) (Hameln, DEXA3.3) and 1× L-Glutamine (Invitrogen) for 6-7d to expand the erythroid progenitor population. Cells were differentiated, over a 30 h period in StemPro medium supplemented with Epo (5 U/mL) (Janssen, PL 00242/029) and transferrin (0.5 mg/mL) (Sigma, T0665) to a late stage of erythropoiesis. Foetal liver material was obtained from mice that are wild-type, DKO (where both MCS-R1 and MCS-R2 are deleted) ^20^, or AMKO (where both α-globin genes are removed)^29^. Mouse ES cell line, E14, was cultured in GMEM (Invitrogen) supplemented 10% (vol/vol) FBS (Gibco®, 10270) and LIF and grown in gelatinised flasks. C127, a mouse mammary epithelial cell line, was cultured in DMEM (Invitrogen) supplemented with 10% (vol/vol) FBS (Sigma), × penicillin/streptomycin (Invitrogen) and 1× L-glutamine (Invitrogen). MEL, the mouse erythroleukamia cell line, was cultured in RPMI (Invitrogen) supplemented with 10% (vol/vol) FBS (Sigma), 1× penicillin/streptomycin (Invitrogen) and 1× L-glutamine (Invitrogen). Mouse embryonic fibroblasts (MEF) were cultured in DMEM (Invitrogen) supplemented with 15% (vol/vol) FBS (Gibco®), 1× penicillin/streptomycin (Invitrogen), 1× L-glutamine (Invitrogen) and 1× NEAA (Invitrogen). All cells were incubated at 37°C in a humidified 5% (vol/vol) CO_2_ incubator. None of the cell lines used here are found in the database of commonly misidentified cell lines that is maintained by ICLAC and NCBI Biosample.

### Fluorescence activated cell sorting

Defined cell populations were obtained as follows; expanded MFL cells were depleted of differentiated erythroid Ter119+ve cells by staining with Ter119 antibody (Becton Dickinson, 553673) and separation using MACS column (Miltenyi Biotec Ltd). Ter119-ve cells were then stained and sorted for CD44 (Becton Dickinson, 561862) and cell size (Fig. 1a). This gave an early erythroid progenitor population. Following 30 h culturing in differentiation medium, intermediate erythroblasts were obtained. Differentiation status was monitored by cytospin, and level of α-globin nascent transcript was assessed by RT-qPCR (Fig. 1b).

### Reverse Transcription qPCR (RT-qPCR)

Isolation of total RNA was performed by lysing 10^7^ cells in TRI reagent (Sigma), according to the manufacturer’s instructions. To remove genomic DNA from RNA samples, samples were treated with TURBO^TM^ DNase according to manufacturer’s protocol (Invitrogen, AM2238). To assess relative changes in gene expression by qPCR, 1 μg of total RNA was used for cDNA synthesis using Superscript^TM^ II reverse transcriptase (Invitrogen, 18064014). Quantification of mRNA levels was performed using SYBR^®^ Green Real Time PCR master mix according to manufacturers instructions (Applied Biosystems, 4309155). The relative standard curve method was used for relative quantitation of RNA abundance.

### Next-generation Capture-C (NG Capture-C)

Performed as previously described^10^. Material was obtained from mES E14 cells and MFL cells (0h and 30h) from WT and AMKO. Briefly, 3C libraries were generated using standard methods similar to the protocol for *in situ* Hi-C. Before oligonucleotide capture, 3C libraries were sonicated to a fragment size of 200 bp and Illumina paired-end sequencing adaptors (New England BioLabs, E6040, E7335 and E7500) were added using Herculase II polymerase (Agilent). Samples were indexed, allowing multiple samples to be pooled before oligonucleotide capture using biotinylated DNA oligonucleotides designed for the α-globin genes Hba1/2, the *MCS-R1* and *–R2* regulatory elements, CTCF - 39.5 (Sigma Aldrich) and the five FISH probe sites Ex, E, A, C, Cx (ATDBio Ltd). The first hybridisation reaction was scaled up relative to the number of samples included in the reaction to maintain library complexity using Nimblegen SeqCap EZ Hybridization and Wash Kit (Roche, 05634261001). After a 72 h hybridisation step, streptavidin bead pulldown (Invitrogen, 65305) was performed, followed by multiple bead washes using Nimblegen SeqCap EZ Hybridization and Wash Kit (Roche, 05634261001) followed by PCR amplification of the captured material using SeqCap EZ accessory Kit v2 (Roche, 07145594001). A second capture step was performed as above, with the exception that it was carried out in a singlevolume reaction. The material was sequenced using the Illumina^®^ MiSeq platform with 150-bp paired-end reads. Data were analyzed using scripts available at https://bitbucket.org/telenius/CCseqBasic and R was used to normalize data and generate differential tracks.

### Probes and nick-translation labelling

For FISH; plasmid probes used were pEx (mm9; chr11 ; 32129812-32136918), pE (mm9; chr11; 32146280-32153457), pA (mm9; chr11; 32201016-32208529), pC (mm9; chr11; 32251235-32258747), pCX (mm9; chr11; 3227598632282385). Probes were constructed in the pBlueScript plasmid by subcloning regions from mouse BAC RP23-469I8 and BAC RP24-278E18 by λ-red-mediated recombination^30^. Mouse BACs were as follows; F1 (RDB 4214 MSMg01-530C17), COMP (RDB 4214 MSMg01-276J20) engineered by λ-red-mediated recombination to give a final insert size covering mm9; chr11 ; 32137046-32200781, and F2 (RP24-278E18). All BACs were obtained from RIKEN^31^ and BACPAC Resources Center (Children’s Hospital Oakland Research Institute; http:/bacpac.chori.org). FISH probes were labeled by nick translation as previously described^32^. Probes were directly or indirectly labeled by nick translation using Cy3-dUTP (GE Healthcare) and digoxygenin-11-dUTP (Roche).

### RASER-FISH

The small size of the locus requires optimal preservation of 3D nuclear structure. However, conventional FISH requires heat denaturation disrupting fine details of chromatin structure below 1 Mb^33,34^. Here we have successfully adapted the principle of chromosome orientation FISH (CO-FISH)^35^, to non-repetitive genomic loci. The resulting RASER (resolution after single-strand exonuclease resection)-FISH method maintains nuclear fine-scale structure by replacing heat denaturation with exonuclease digestion, and is suitable for high- and super-resolution imaging analysis. Line profiles across DAPI-stained nuclei after three separate treatments (immunofluorescence (IF), our standard 3D-FISH and RASER-FISH) indicated a loss of structure in the 3D-FISH nuclei that is not observed in the RASER-FISH nuclei when compared to IF only (Extended Data Fig. 8). When comparing RASER-FISH to 3D-FISH, hybridisation efficiency was similar for the two techniques (>90%), suggesting that exonuclease digestion around the alpha globin locus is extensive by this method. Briefly, cells were labelled overnight with BrdU/BrdC mix (3:1) at final conc of 10 μM. Cells were fixed in 4% PFA (vol/vol) for 15 min and permeabilized in 0.2% Triton X-100 (vol/vol) for 10 min. Cells were then stained with Hoechst 33258 (0.5 μg/mL in PBS), exposed to 254 nm wavelength UV light for 15 min, then treated with Exonuclease III (NEB) at final conc 5 U/μL at 37°C for 15 min. Labelled probes (100 ng each) were denatured in hybridization mix at 90°C for 5 min, BACs were preannealed at 37°C for 20 min. Coverslips were hybridized overnight with prepared probes at 37°C. After hybridization, coverslips were washed for 30 min twice in 2× SSC at 37°C, once in 1× SSC at RT. Coverslips were blocked in 3% BSA (wt/vol) and digoxigenin was detected with sheep anti-digoxigenin FITC (Roche, 11207741910) followed by rabbit anti–sheep FITC (Vector Laboratories, FI-6000). Coverslips were stained with DAPI (0.5 μg/mL in PBS), washed with PBS and mounted in Slowfade^®^ Diamond mountant for standard widefield imaging (Molecular Probes^®^) or in Vectashield for STED imaging (Vector Laboratories).

### Standard 3D DNA-FISH

3D DNA-FISH was performed as described previously^36^. In brief, cells were fixed in 4% PFA (vol/vol) for 15 min and permeabilized in 0.2% Triton X-100 (vol/vol) for 10 min. Cells were denatured in 3.5 N HCl for 20 min and neutralized in ice-cold PBS. Probes were prepared as in the previous section, and coverslips were hybridized overnight at 37°C. Cells were washed and blocked, probes were detected and coverslips were mounted as in the previous section.

### Tolerance

Pools of oligonucleotide probes were designed consisting of 30 nt tiling 6 kb of the MCSR2 region, avoiding large repeats, with 30 nt gaps between probes (80 oligonucleotides in total). The probes were synthesised with 5’-amino groups using standard phosphoramidite chemistry (ATDBio Ltd). After purification by gel filtration, the probes were labelled in pools covering 1 kb with either digoxigenin NHS ester or Cy3 NHS ester, to give a 6 kb probe with alternating 1 kb regions of Cy3 or digoxigenin. Conditions for labelling: 1 mM oligonucleotides, 10 mM NHS ester (added as 0.1 M solution in DMSO), 0.5 M sodium carbonate buffer pH 8.5, shaken at 55 °C for 5 h and purified by gel filtration followed by RP-HPLC eluting with a 0.1 M TEAA/MeCN gradient. Fractions containing the products were combined, dried, desalted by gel filtration and lyophilised. MEL cells were fixed on coverslips and prepared according to the RASER-FISH protocol. The pooled probes were resuspended in water at 100 ng/ μL. 1 μL of the labelled oligonucleotide mixture was added to 5 μL hybridisation buffer (Kreatech) and 5 μL 2X SSC. The probe mixture was denatured at 95°C for 5 min, placed on ice, then applied to the coverslip. The coverslips were hybridized at 37°C overnight, then washed, detected and mounted as previously described.

### Imaging Equipment and settings

Widefield fluorescence imaging was performed at 20°C on a DeltaVision Elite system (Applied Precision) equipped with a 100×/1.40 NA UPLSAPO oil immersion objective (Olympus), a CoolSnap HQ2 CCD camera (Photometrics), DAPI (excitation 390/18; emission 435/40), FITC (excitation 475/28; emission 525/45) and TRITC (excitation 542/27; emission 593/45) filters. 12-bit image stacks were acquired with a z-step of 150 nm giving a voxel size of 64.5 nm × 64.5 nm × 150 nm. Image restoration was carried out using Huygens deconvolution Classic Maximum Likelihood Estimation (Scientific Volume Imaging B.V.), STED images were acquired at 20°C on a Leica TCS SP8 3X Gated STED (Leica Microsystems), equipped with a pulsed supercontinuum white light excitation laser at 80Mhz (NKT), and two continuous wavelength STED lasers at 592 nm and 660 nm. HyD detectors were used in gated mode (1.5-6ns for 592 depletion and 0.5-8.5ns for 660 depletion) A sequential imaging mode was set employing first the 660 nm STED laser, and then the 592 nm STED laser to give a final voxel size of 31.9 nm × 31.9 nm × 110 nm in the image shown (Figure 3b), which was minimally smoothed by performing a Gaussian blur of 0.75 pixel radius in ImageJ (https://imagej.net/).

### Image Analysis

Measurements of either distance or volume were made using in-house scripts *(Note; available via Github upon manuscript acceptance)* in ImageJ. As a preprocessing step image regions are chromatically corrected to align the green and the red channel images. The parameters for the chromatic correction were calculated through taking measurements from images of 0.1 μm TetraSpeck^®^ (Molecular Probes^®^) and calculating the apparent offset between images in each colour channel. For both distance and volume measurement scripts, signals were manually selected by a single click whereupon a 20 × 20 pixel and 7 × z-step sub-volume was generated centred on the identified location (Extended Data Fig. 9a). In each selected region, thresholding was applied to segment the foci. Firstly the image region was saturated beyond the top 96.5 % intensity level, to reduce the effect of noisy pixels, and then the threshold was calculated as being 90 % of the maximum intensity value of the processed image. This was repeated for both green and red channels and was found to accurately segment the foci from background. Once segmented the inter-centroid 3-D distance calculation was made between the centroids in 3-D and output along with a .png image for visual inspection (Extended Data Fig. 9a). For the volume analysis, the segmented volume for foci was integrated and converted into μm^3^ units and output for each signal. We validated any increase in volume between MFL 0h and 30h by taking volume measurements of fluorescently labelled 500 nm diameter Tetraspeck^®^ beads (Molecular Probes^®^) incorporated into the mountant where we found the bead volume measurements equivalent at the two time points (data not shown). Correlation of the positioning of paired FISH probes was assessed by Pearson co-efficient of correlation analysis and was performed on the 20x20x7 raw intensity signals from each channel. Line profile analysis was performed using the Plot Profile function in Fiji^37^. We made initial comparisons between z-steps of 100 nm, 150 nm and 200 nm to assess any effect on inter-centroid 3-D distance measurements (Extended Data Fig. 9b) and established the tolerance of the inter-centroid distances produced by the analysis pipeline to be 53 nm (Extended Data Fig. 9c).

### Statistics and reproducibility

Statistical analysis was carried out with Graphpad Prism (version 7.0c) unless otherwise indicated. Gene expression experiments were performed on three biological replicates (standard deviation (s.d.) is shown). All NG Capture-C experiments were performed on three biological replicates with the exception of WT and AMKO capture from R1/R2 which were each derived from one sample. The standard deviation of 250 bp bins was calculated in R and visualized to illustrate the reproducibility of this chromatin interaction analysis. All graphs showing FISH signal inter-distance data display median values with interquartile range with the exception of Extended Data Fig. 9c, which show mean values with s.d.. All volume analyses are displayed as cumulative frequency plots where the bins were in voxel sized increments. The statistical significance of differences in the range of distance measurements and volume measurements were derived as two-tailed by the Kruskal-Wallis test with Dunn’s multiple comparisons. *P* values are represented as \**P* <0.05; \*\**P* <0.01; \*\*\**P* <0.001; \*\*\*\**P* <0.0001.

## Acknowledgements

We thank S. Butler for tissue culture support, J. Sloane-Stanley and J. Sharp for mouse breeding and foetal liver provision, K. Clark and C. Waugh of the Flow Cytometry Facility for FACS analysis, E. Repapi for advice on statistical analysis, E. Garcia for advice on STED imaging, C. Harrold and J. Davies for analysis of NG Capture-C data, T. Brown for oligonucleotide synthesis support. This work was supported by the Medical Research Council (MC_UU_12009 to V.J.B., D.H., J.H. and MR/N00969X/1 to J.H.) and Wellcome Trust (106130/Z/14/Z to DH). Further support came from grants to the Wolfson Imaging Centre Oxford (Wolfson Foundation 18272, joint MRC/BBSRC/EPSRC MR/K015777X/1, Wellcome Trust Multi-User Equipment 104924/Z/14/Z) and the WIMM FACS Core Facility (NIHR Oxford BRC and John Fell Fund (131/030 and 101/517), the EPA fund (CF182 and CF170) and by the WIMM Strategic Alliance awards G0902418 and MC_UU_12025).

## Data availability

Capture-C data generated for this study have been deposited in the Gene Expression Omnibus (GEO) under accession code *(in process of submission)*. All images files are archived in OMERO and can be made available upon request. Analysis scripts for distance and volume measurements are available at https://github.com/dwaithe/focimeasurements *(Note; this will be activated upon manuscript acceptance)*. All other data supporting the findings of this study are available from the corresponding author on reasonable request.

## Author Contributions

V.J.B. and D.R.H. conceived the project. J.M.B. and V.J.B. developed the RASER-FISH technique from Co-FISH and performed the FISH experiments with assistance from I.S. C.B. sub-cloned the FISH probes and S.D.O. synthesised oligonucleotides. C.L. assisted with the imaging and image storage. D.W. wrote scripts for and advised on image analysis. A.M.O. designed analysis of the α-globin domain by NG Capture-C. N.R. and J.T. undertook the NG Capture-C experiments and subsequent analysis respectively. B.G. and M.T.K. developed the erythroblast *ex vivo* differentiation system and B.G. performed the FACS analysis and nascent transcript quantification. V.J.B., J.M.B., J.R.H. and D.R.H. wrote the paper.

## Author Information

The authors declare no competing financial interests. Correspondence and requests for materials should be addressed to V.J.B.

---

**Extended Data Figure 1:**
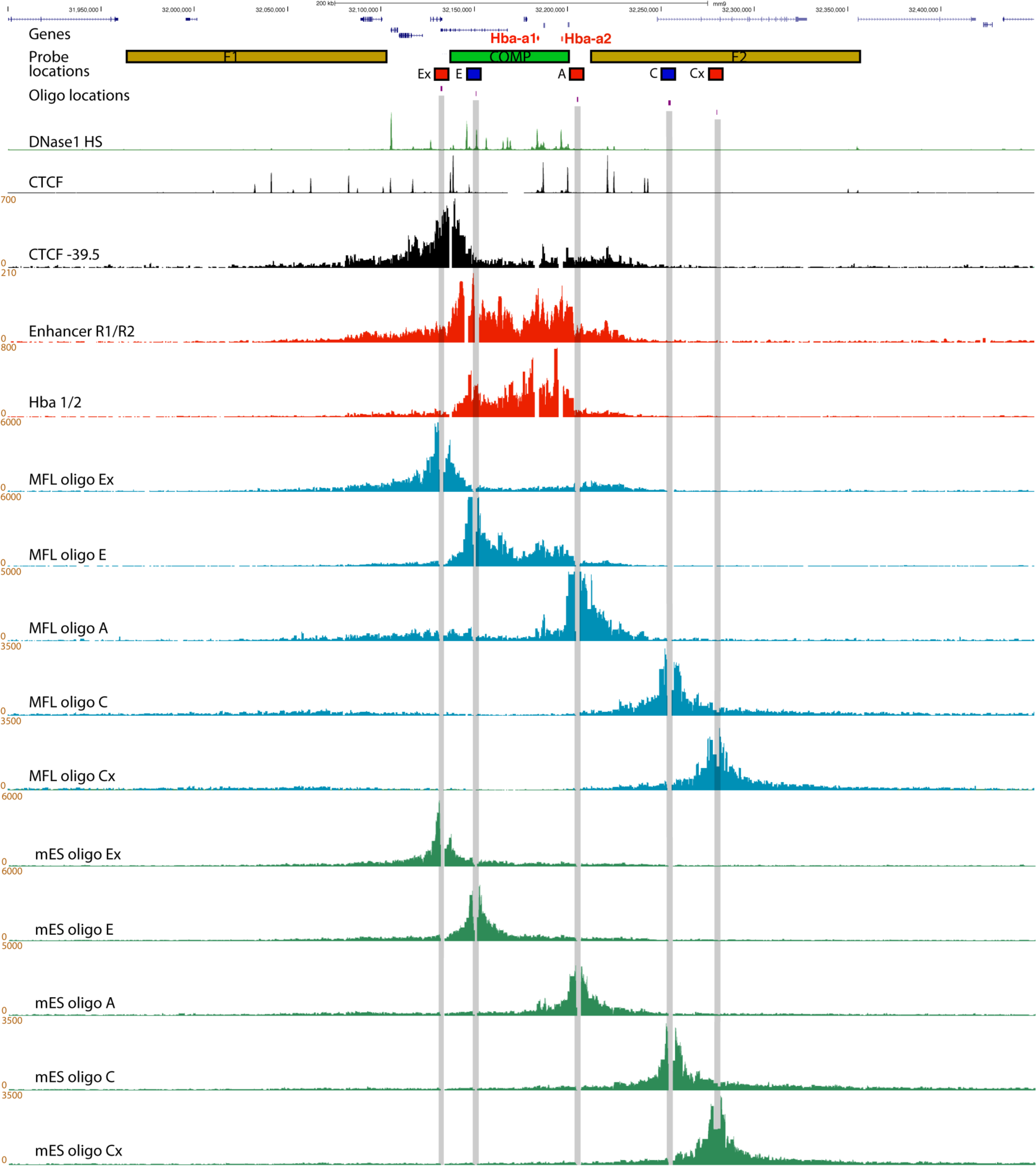
Chromatin interactions detected from FISH probe viewpoints in erythroblasts and mES cells. Hba genes are highlighted in red, followed by the locations of the BAC (F1, COMP, F2) and plasmid (Ex, E, A, C, Cx) FISH probes and the 50mer oligonucleotides used for capture. Underneath are genome browser tracks showing DNase1 hypersensitive sites ’ (green) and CTCF BS (black), then NG Capture-C tracks for MFL 30h using the CTCF −39.5 BS (black), the two major enhancer elements MCS-R1/-R2, and the Hba genes as viewpoints (both red). Below are five NG Capture-C tracks from MFL 30h erythroblasts (blue) depicting interactions from the viewpoints of the five FISH probes, as indicated. Interactions detected by oligo Ex mirror those detected from the CTCF BS −39.5 at the upstream side of the α-globin domain; oligo E detects interactions within the self-interacting domain, matching the Enhancer R1/R2 track; oligo A detects interactions at the opposite side of the domain whilst control oligos C and Cx principally detect proximity interactions. Measurements A to E will therefore reflect a mixture of interactions within and across the domain. Five further tracks (green) depict interactions in E14 mES cells where only proximity interactions are detected.

**Extended Data Figure 2:**
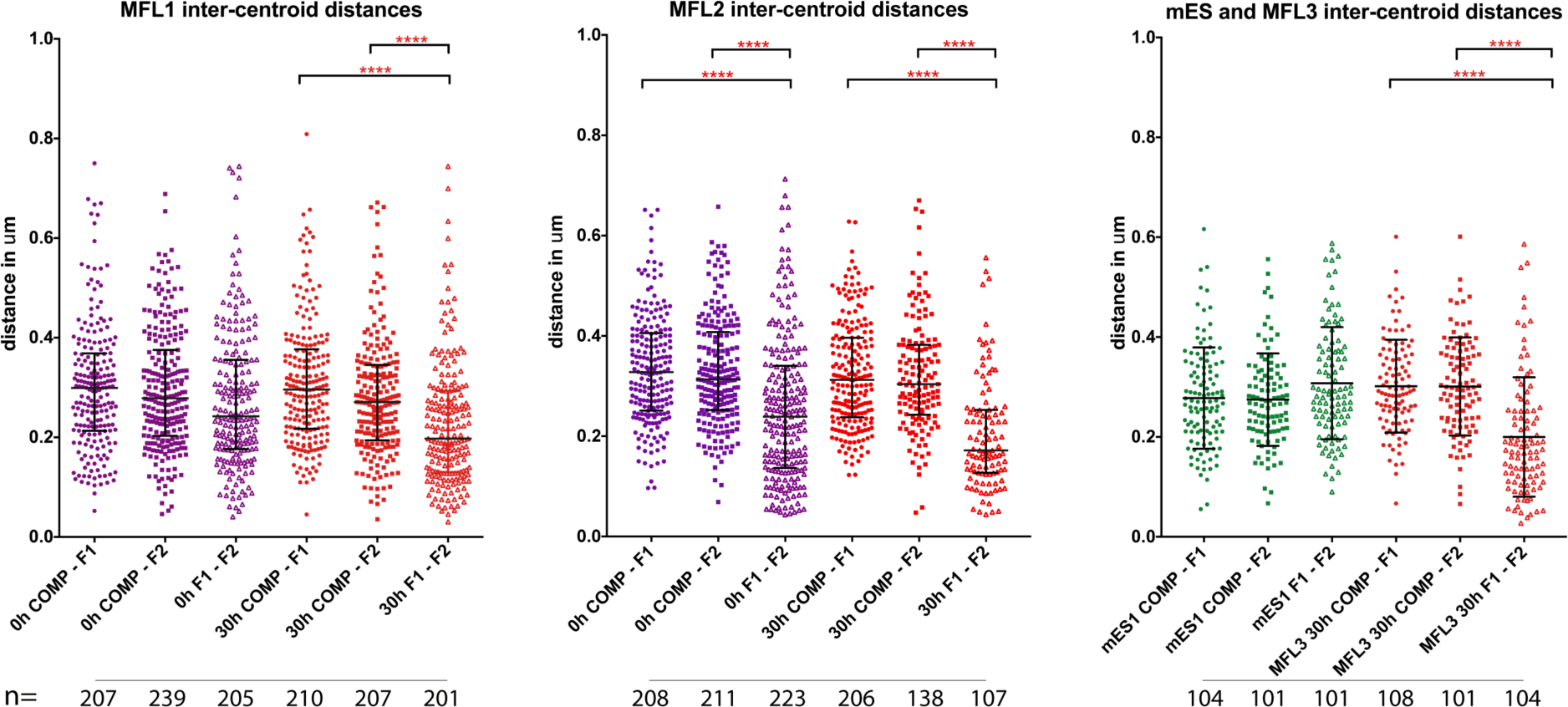
Proximity measurements across a 320 kb region encompassing the α-globin locus in mouse WT cells. Inter-centroid distance measurements between three BAC probe pairs, COMP-F1 (circles), COMP-F2 (squares) and F1-F2 (open triangles) in erythroblasts derived from three independent MFL cultures and in mES cells. MFL1 and MFL2 measurements are given for two time points, 0h and 30h. Each dot represents a single measurement. Error bars indicate median value and interquartile range. Statistical significance of differences in range of measurements, derived by a Kruskal-Wallis test with Dunn’s multiple comparisons, is shown (****p<0.0001).

**Extended Data Figure 3:**
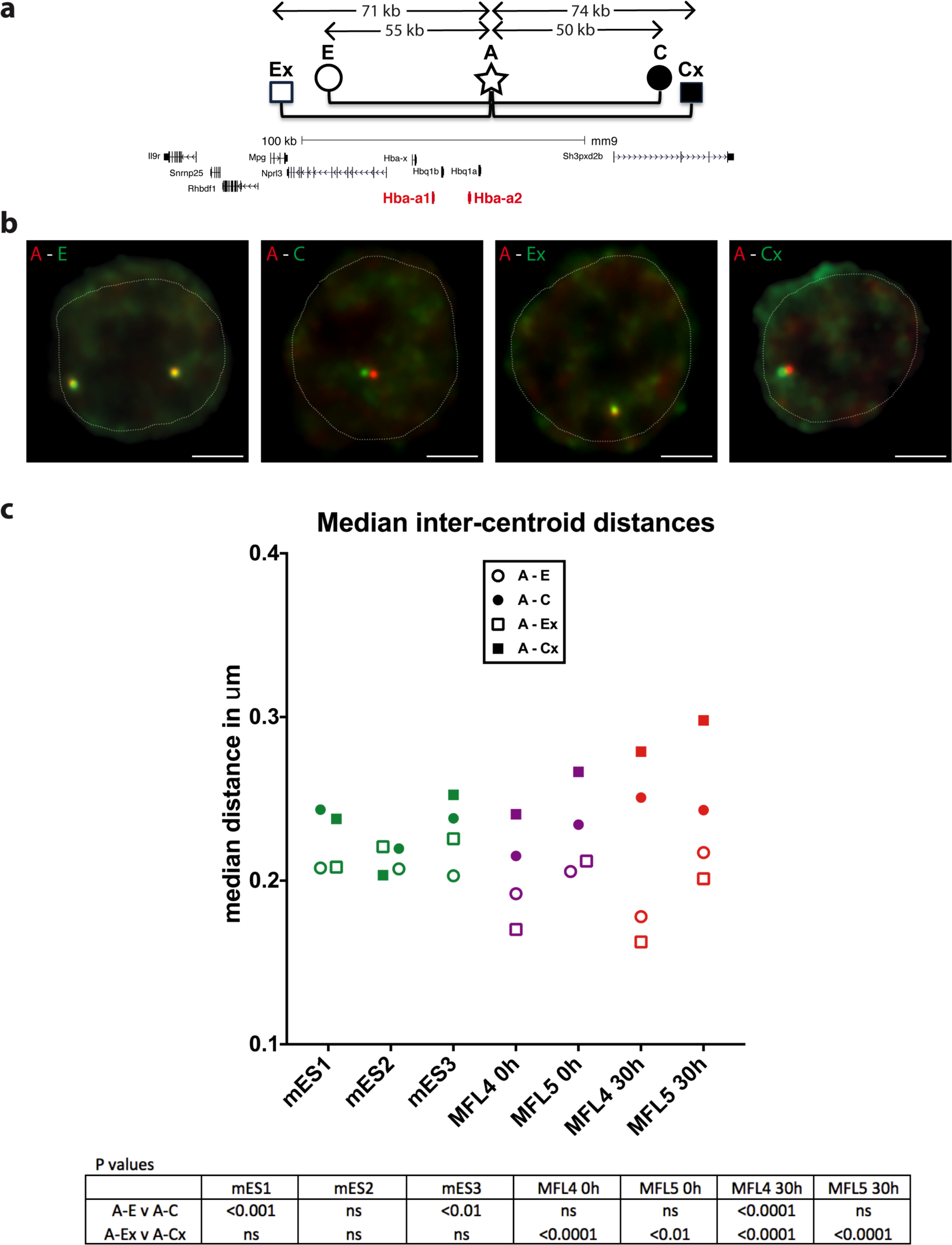
Proximity measurements at the α -globin locus in mouse WT cells. **a,** Gene map with plasmid FISH probe locations showing the pairwise combinations used to measure inter-probe distances. Using probe A as a point of reference, measurements were made to the domain side (E, Ex) of probe A compared to the control non-interacting side (C, Cx). Genomic distances between midpoints of the probe pairs are shown. **b,** Representative images of RASER-FISH hybridisation signals for the four plasmid probe pairs in MFL 30h erythroblasts. White dotted line delineates nuclei. Scale bar 2μm. **c,** Median inter-centroid distances measured between the four probe pairs in three different cell types, mES1-3 (green), MFL4-5 0h (purple) and MFL4-5 30h (red). P values, derived by a Kruskal-Wallis test with Dunn’s multiple comparisons, are shown. See Extended Data Fig. 4 for full data with statistical analyses. At MFL 30h but not mES, the distance between A and Ex is consistently statistically shorter (p<0.0001) than A to Cx.

**Extended Data Figure 4:**
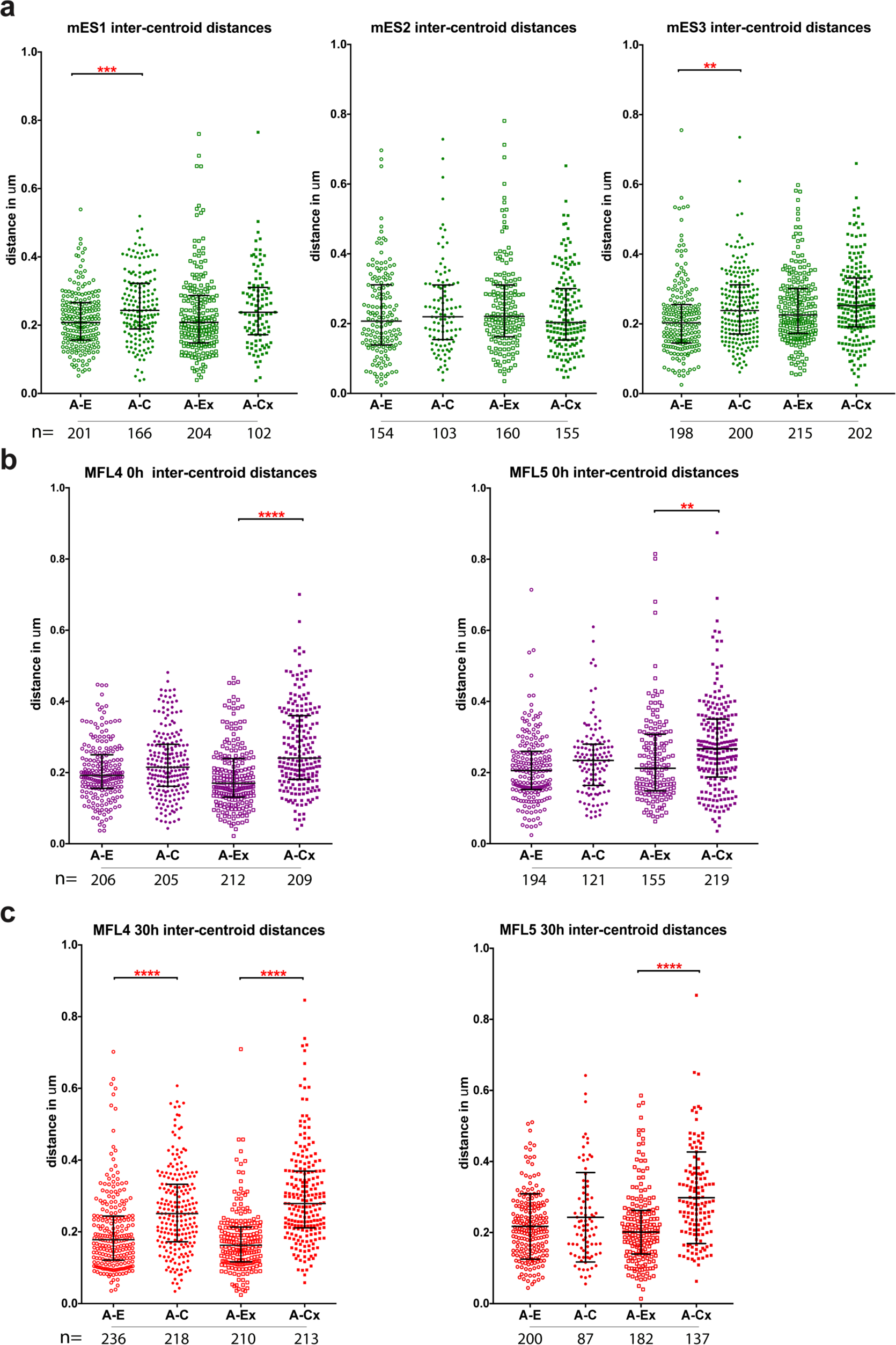
Proximity measurements at the α -globin locus between plasmid probe pairs. **a,** Inter-centroid distances measured between the four probe pairs A-E (open circles), A-C (closed circles), A-Ex (open squares), A-Cx (closed squares) in three mES cell cultures (green). Each dot represents a single measurement. Error bars indicate median value and interquartile range. Statistical significance of differences in range of measurements, derived by a Kruskal-Wallis test with Dunn’s multiple comparisons, is shown. **b,** Inter-centroid distances plotted as above for two MFL cultures harvested at 0h (purple). c, Inter-centroid distances plotted as above for two MFL cultures harvested at 30h (red). ****p< 0.0001; ***p< 0.001; **pm 0.01.

**Extended Data Figure 5:**
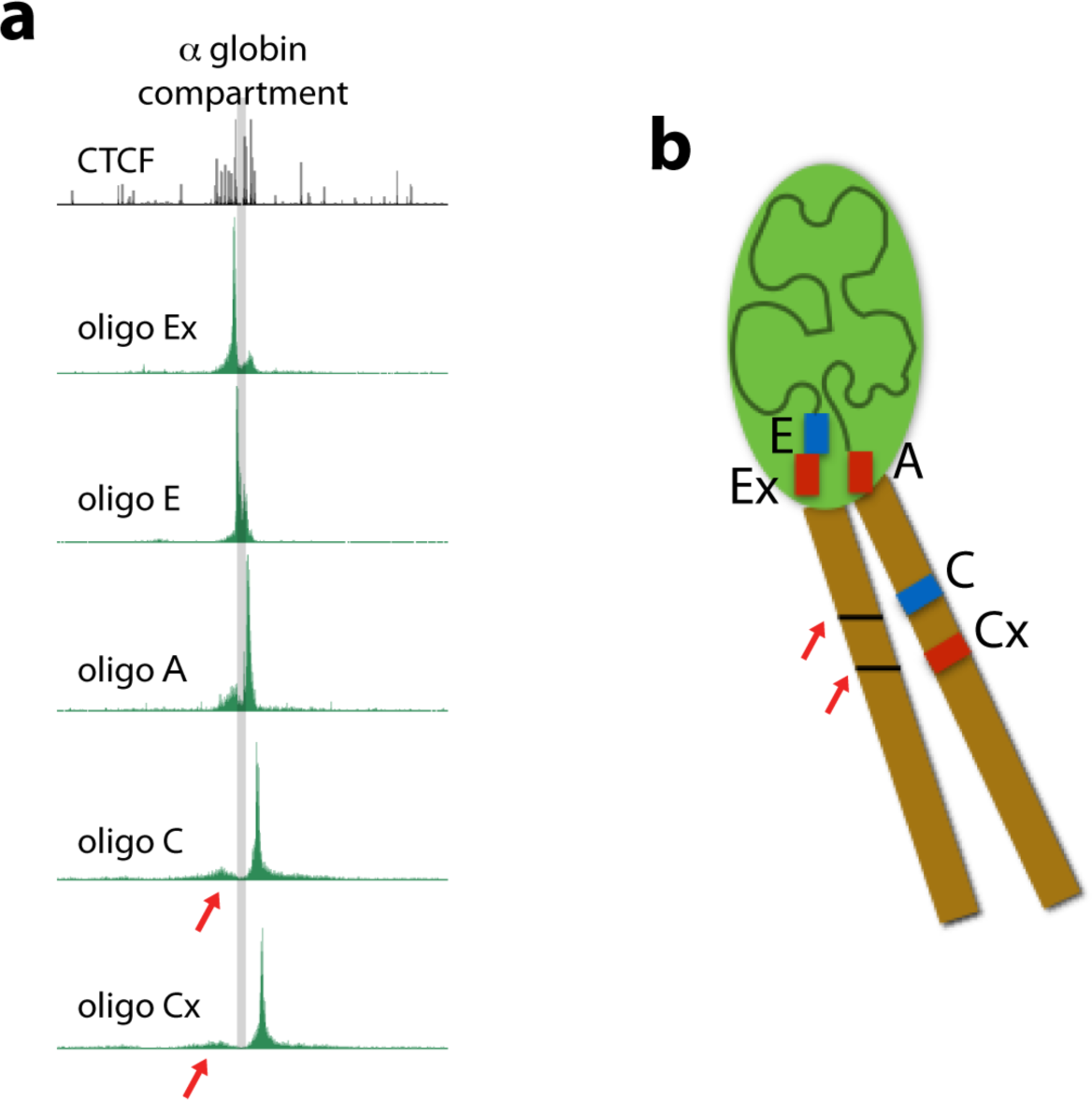
Infrequent interactions occur between chromatin regions encompassing the α -globin self-interacting domain. **a,** A larger scale view of NG Capture-C tracks for MFL30h from plasmid probe viewpoints as presented in Extended Data Fig. 1. The extent of the α-globin domain is defined by the pale grey bar. Outlying interactions detected by oligo C and Cx to a region devoid of genes or erythroid-specific accessibility are indicated by red arrows. Careful examination indicates that C interacts rather more frequently and with a region that is slightly closer than Cx. Such interactions are consistent with the development of a distinct domain that affects the positioning of the flanking regions. **b,** Model of the domain showing that the structure created by the self-interacting domain can lead to more frequent interactions between surrounding chromatin.

**Extended Data Figure 6:**
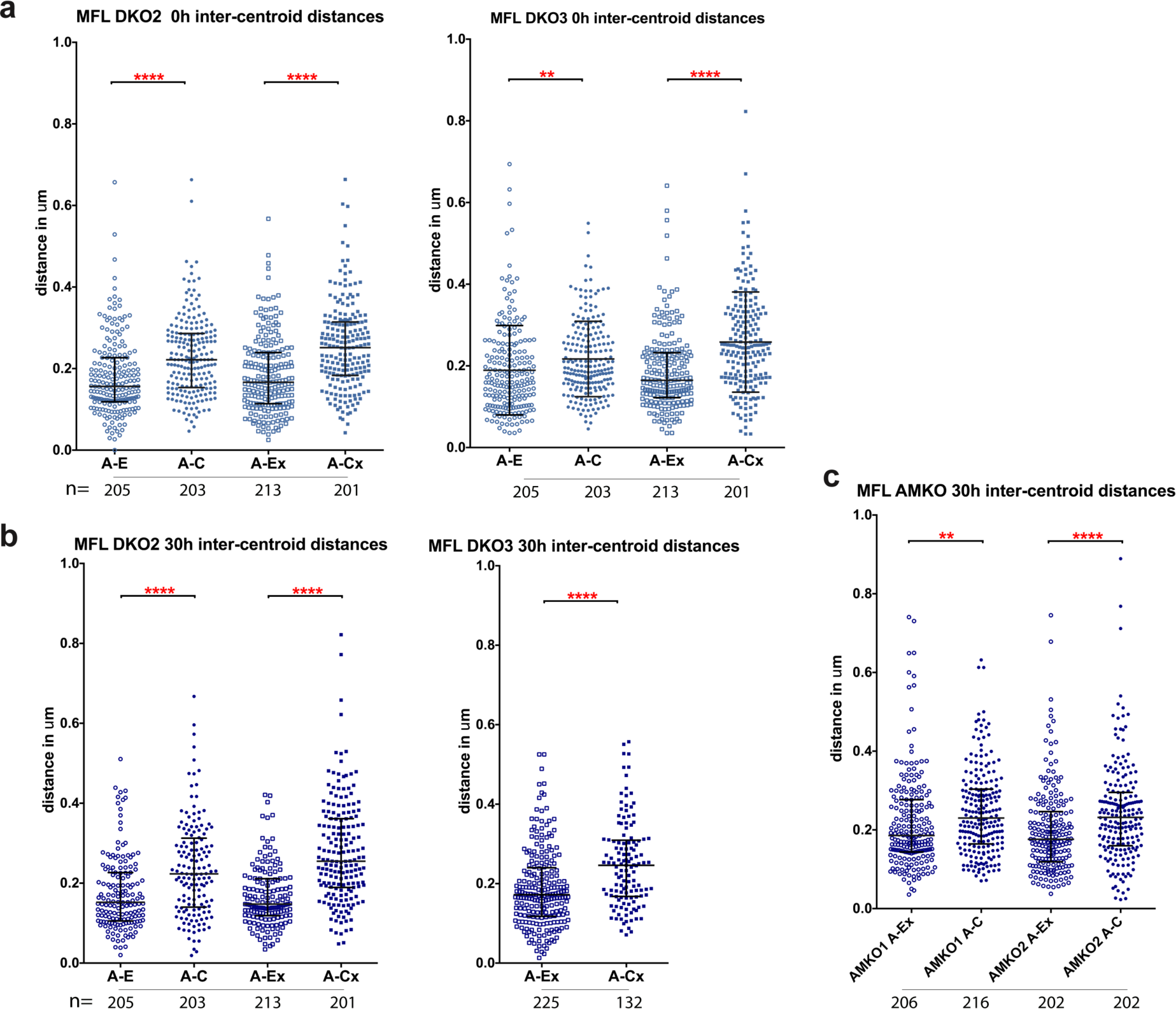
Proximity measurements at the α-globin locus between plasmid probe pairs in knockout mouse lines. **a,** Inter-centroid distances measured between the four probe pairs A-E (open circles), A-C (closed circles), A-Ex (open squares), A-Cx (closed squares) in two MFL DKO erythroblast cultures at 0h (light blue). Each dot represents a single measurement. Error bars indicate median value and interquartile range. Statistical significance of differences in range of measurements, derived by a Kruskal-Wallis test with Dunn’s multiple comparisons, is shown. b, Inter-centroid distances plotted as above for two MFL DKO cultures harvested at 30h (dark blue). c, Inter-centroid distances plotted as above for two MFL AMKO cultures harvested at 30h (dark blue). ****p< 0.0001; ***p< 0.001; **p< 0.01.

**Extended Data Figure 7:**
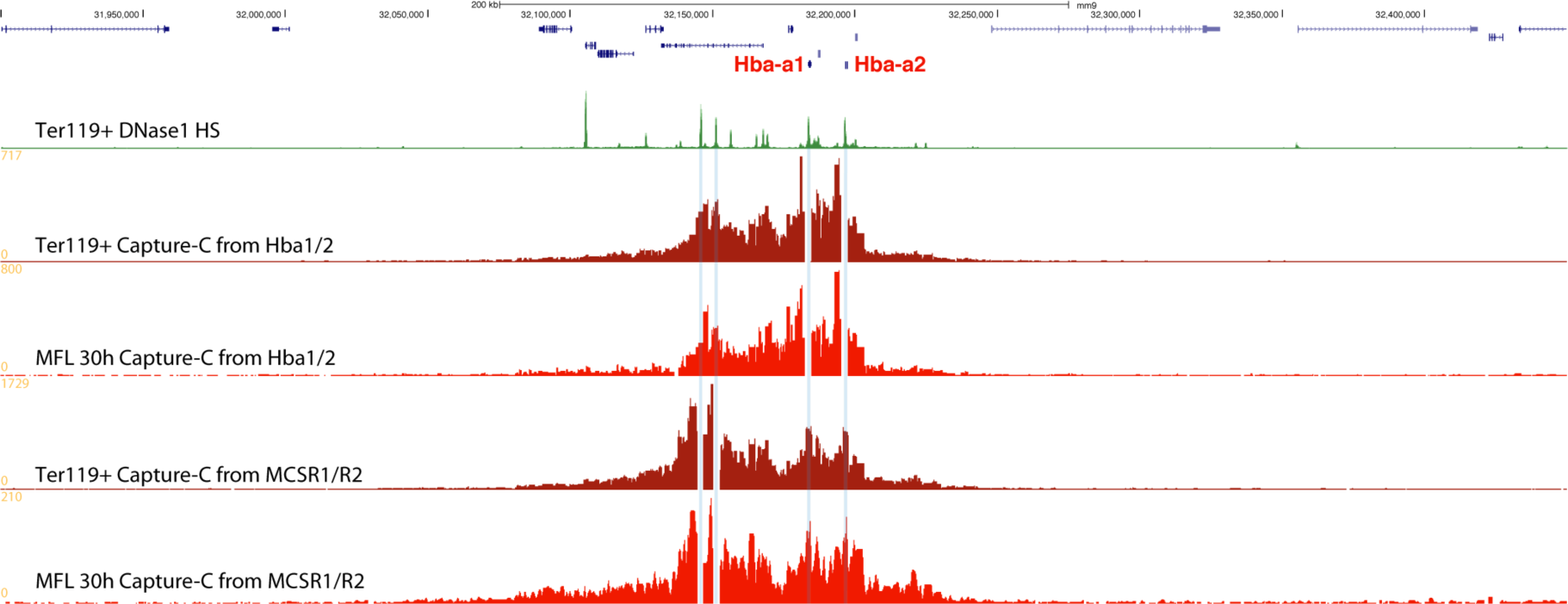
Erythroblasts derived from adult spleen or foetal liver form the same domain of chromatin interactions at the α -globin locus. Layout as for Fig. 1c. Capture-C tracks from Hba1/2 and MCSR1/R2 viewpoints show the same pattern of chromatin interactions between erythroblasts derived from adult spleen (Ter119+ dark red) and foetal liver (MFL 30h red).

**Extended Data Figure 8:**
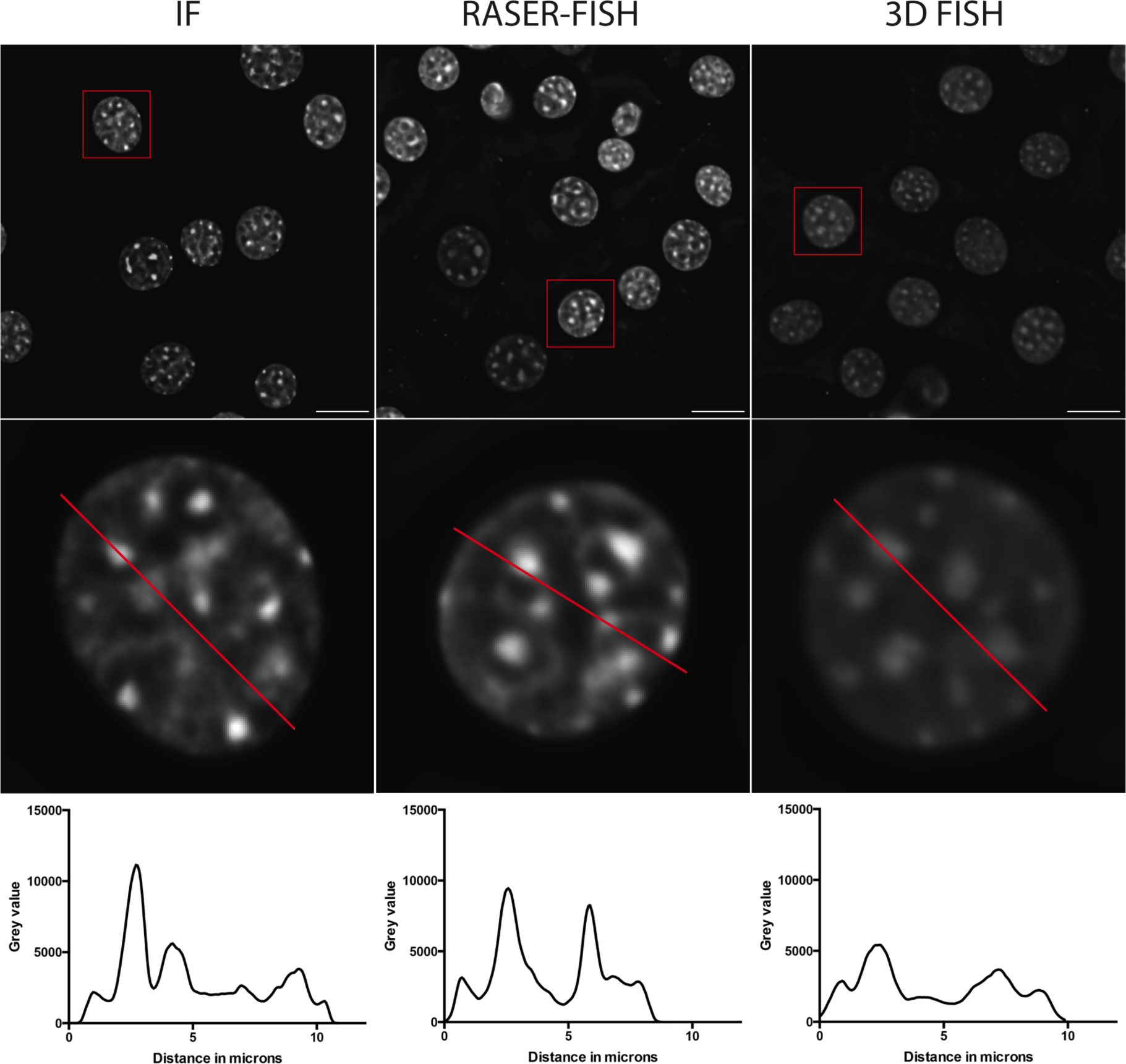
The RASER-FISH technique. Example C127 DAPI-stained nuclei (top) after fixation and immunofluorescence only (left), RASER-FISH (middle) or 3D-FISH (right). Bar = 10μm. Red box indicates selected nucleus; red line across these nuclei (middle) indicates position of line profiles (bottom) indicating fluorescence intensity across the matching nucleus, reduced after 3D-FISH.

**Extended Data Figure 9:**
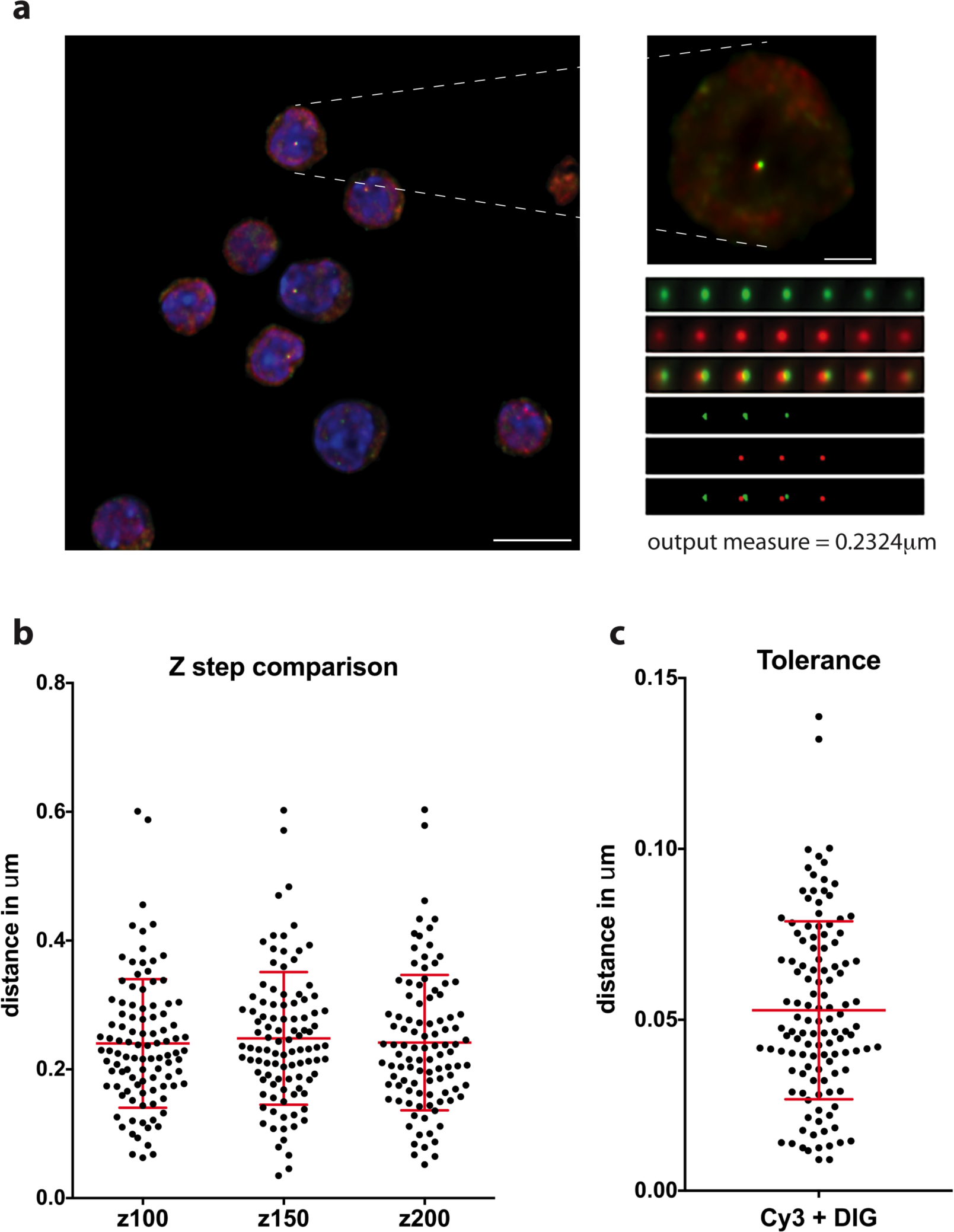
Image capture and analysis. **a,** Example field of capture after deconvolution (left) with selected nucleus (right), above the sub-stack of 7 Z steps generated from the nuclear signal. The signal is thresholded at 80% of maximum fluorescence then distance between signal centroids is calculated in three dimensions after chromatic shift correction. Scale bars 10 μm and 2 μm. **b,** A-C distance measurements from the same signal pairs were taken after collection of the image stacks at three different Z steps; 100, 150 and 200 nm. There is no difference between the spread of data or mean values for the three data sets. n=101. **c,** The tolerance of a FISH experiment represents the distance that can be measured between different fluorescent labels to the same probe. Here we hybridised two pools of oligos for MCS-R2 directly labelled with Cy3 and digoxigenin and find a mean tolerance of 53 nm, well below measurements across the α-globin locus. n=121.

